# Engineering Cell-Permeable Proteins through Insertion of Cell-Penetrating Motifs into Surface Loops

**DOI:** 10.1101/2020.01.03.894543

**Authors:** Kuangyu Chen, Dehua Pei

## Abstract

Effective delivery of proteins into the cytosol and nucleus of mammalian cells would open the door to a wide range of applications including treatment of many currently intractable diseases. However, despite great efforts from numerous investigators and the development of a variety of innovative methods, effective protein delivery in a clinical setting is yet to be accomplished. Herein we report a potentially general approach to engineering cell-permeable proteins by genetically grafting a short cell-penetrating peptide to an exposed loop region of a protein of interest. The grafted peptide is conformationally constrained by the protein structure, sharing the structural features of cyclic cell-penetrating peptides and exhibiting enhanced proteolytic stability and cellular entry efficiency. Insertion of an amphipathic motif, Arg-Arg-Arg-Arg-Trp-Trp-Trp, into the loop regions of enhanced green fluorescent protein (EGFP), protein-tyrosine phosphatase 1B (PTP1B), and purine nucleoside phosphorylase (PNP) rendered all three proteins cell-permeable and biologically active in cellular assays. When added into growth medium, the engineered PTP1B dose-dependently reduced the phosphotyrosine levels of intracellular proteins, while the modified PNP protected PNP-deficient mouse T lymphocytes (NSU-1) against toxic levels of deoxyguanosine, providing a potential enzyme replacement therapy for a rare genetic disease.

## INTRODUCTION

Cell-permeable proteins would provide powerful research tools as well as novel therapeutics against many of the currently undruggable targets, such as intracellular protein-protein interactions and missing/defective proteins caused by genetic mutations. Unfortunately, because of their large sizes and hydrophilic nature, proteins are generally impermeable to the cell membrane. Interestingly, a small number of naturally occurring proteins including the trans-activator of transcription (Tat) protein of HIV,^1,2^ the homeobox protein Antennapedia of *Drosophila*,^3^ bacterial toxins,^4^ and human protein α-synuclein^5^ have demonstrated the capacity of entering the cytosol of mammalian cells. It was later found that short peptides derived from some of these proteins, e.g., Tat^47–57^ from the HIV Tat protein^6^ and penetratin from Antennapedia^7^, are sufficient to mediate the internalization of these proteins. Since these initial discoveries in the 1990’s, hundreds of other short peptides (typically 5-30 residues) of both natural and synthetic origins have been found to have similar cellular entry capabilities.^8–10^ These peptides are collectively termed as “cell-penetrating peptides (CPPs)”. The CPPs have been applied to deliver a wide variety of cargoes including small molecule drugs, peptides, proteins, nucleic acids, and nanoparticles into mammalian cells.^8–10^ For example, several CPPs have been genetically fused to the N- or C-terminus of an otherwise impermeable protein to render the latter cell-permeable.^11–14^ An attractive feature of CPP-mediated protein delivery is that CPPs are relatively tolerant to the physicochemical properties of the cargo, which are highly diverse among different proteins. However, linear CPPs generally have low cytosolic delivery efficiencies (<5%).^15,16^ It is now generally accepted that CPP-protein conjugates enter mammalian cells predominantly by endocytic mechanisms and become entrapped inside the endosome and lysosome, with only a very small fraction reaching the cytosol, which is the intended destination for most applications. Another major limitation of CPP-mediated protein delivery is the poor proteolytic stability of the CPP-protein conjugates. CPPs genetically fused to the N- or C-terminus of a cargo protein consist of proteinogenic amino acids and are generally unstructured. Consequently, the fusion protein is highly susceptible to proteolytic degradation in vivo (e.g., during circulation), resulting in rapid loss of the CPP sequence. In fact, some of the fusion proteins are so unstable that they cannot be isolated from bacterial expression systems in their intact forms.^14^

Several other approaches are also being pursued to deliver recombinant proteins into mammalian cells, including physical methods,^17^ fusion with bacterial toxins,^18^ surface modification,^19–21^ complexation with cationic peptides and polymers,^22–25^ and encapsulation into nanoparticles and liposomes.^26,27^ Each of these approaches has its advantages but also faces unique challenges. Protein delivery by physical methods is generally incompatible with clinical applications, except for topical administration. Fusion of protein cargoes with a bacterial toxin allows for endocytic uptake specifically into target cells/tissues and efficient endosomal escape, but is highly immunogenic and has limited capacity (in terms of the number of cargo molecules delivered into each cell). Supercharging proteins by replacing their non-essential surface residues with positively charged amino acids results in highly efficient endocytic uptake, but the endocytosed proteins remain entrapped inside the endosomal/lysosomal compartments, with little cytosolic exposure.^16^ Complexation with cationic polymers is possible for negatively charged proteins but less effective for neutral and positively charged ones. Finally, protein delivery by encapsulation into nanoparticles or liposomes may prove challenging for tissues outside the liver, spleen, and kidney.^28^ Clearly, there remains an urgent need for alternative approaches that more effectively deliver proteins into the mammalian cytosol.

Others^29–31^ and we^32,33^ recently discovered that cyclization of CPPs enhances their cellular entry efficiencies. An added benefit of peptide cyclization is the greatly improved proteolytic stability. In particular, we reported a family of small amphipathic cyclic peptides as exceptionally active CPPs.^34^ The cyclic CPPs have been applied to efficiently deliver a variety of drug modalities, including proteins^33,35,36^ and nucleic acids,^37,38^ into the mammalian cytosol in vitro and in vivo. However, a drawback of the cyclic CPPs, which contain non-proteinogenic amino acids, is that they must be conjugated to a protein cargo post-translationally. Since site-specific modification of proteins remains a significant challenge,^39^ especially on an industrial scale, it is highly desirable to integrate the cyclic CPP sequence into the structure of a cargo protein, so that the CPP-protein conjugate can be produced recombinantly. In this study, we show that CPP motifs can indeed be genetically inserted into the loop regions of a cargo protein to mimic the conformations of cyclic CPPs, resulting in proteolytically stable proteins and their effective delivery into the cytosol of mammalian cells.

## RESULTS AND DISCUSSION

### General Design Considerations

We chose to fuse CPP motifs to the loop regions (instead of the N- or C-terminus) of cargo proteins for several reasons. First, insertion of a short CPP peptide into a surface loop or replacement of the original loop sequence with a CPP should constrain the CPP sequence into a “cyclic” conformation, greatly enhancing its proteolytic stability. Second, the “cyclic” conformation of a loop-embedded CPP may mimic that of a cyclic CPP and enhance its cellular entry efficiency. Third, previous studies have shown that insertion of proper peptide sequences into surface loops of a protein often causes only minor destabilization of the protein structure.^40^

Another important consideration is the CPP sequence. CPPs are thought to escape the endosome by binding to the intraluminal membrane and inducing CPP-enriched lipid domains to bud off the endosomal membrane as tiny vesicles, which then disintegrate into amorphous lipid/CPP aggregates inside the cytoplasm.^34^ Amphipathic CPPs likely facilitate endosomal escape by stabilizing the budding neck structure, which features simultaneous positive and negative membrane curvatures in orthogonal directions (or negative Gaussian curvature).^41^ It is hypothesized that the hydrophobic group(s) may insert into the lipid bilayer to generate positive curvature, while the arginine residues bind to and bring phospholipid head groups together, inducing negative membrane curvature. In addition, the most active cyclic CPPs [e.g., cyclo(Phe-phe-Nal-Arg-arg-Arg-arg-Gln),^34^ where phe is D-phenylalanine, Nal is L-naphthylalanine (Nal), and arg is D-arginine] contain D-as well as L-amino acids at roughly alternating positions. Recombinant production of proteins necessitates the use of genetically encoded amino acids. Although some of the non-proteinogenic amino acids in cyclic CPPs (e.g., Nal) may be incorporated into proteins by expanding the genetic code,^42^ we decided to use only the 20 proteinogenic amino acids as building blocks in this study for technical simplicity. We chemically synthesized a small panel of amphipathic peptides containing 3 or 4 arginine and 2 or 3 phenylalanine and/or tryptophan residues and cyclized them via a disulfide bond between two cysteine residues flanking the CPP sequence. The peptides were labeled with a pH-sensitive dye, naphthofluorescein (NF), and HeLa cells treated with the peptides were analyzed by flow cytometry. Because NF is nonfluorescent inside the acidic environments of the endosome and lysosome, the cellular fluorescence as determined by flow cytometry reflects the cytosolic entry efficiency of the peptide.^43^ Among the peptides tested, cyclo(CWWWRRRC) showed the highest cytosolic entry efficiency and was selected for protein delivery experiments.

### Intracellular Delivery of EGFP by Inserting CPPs into a Surface Loop

We first tested the loop insertion strategy on enhanced green fluorescent protein (EGFP), whose intrinsic fluorescence facilitates the identification of properly folded mutants as well as the assessment of cellular entry efficiency. Loop 9 of EGFP (amino acids 171-176) was previously shown to be highly tolerant to peptide insertion.^44^ We therefore inserted the CPP motif WWWRRR between Asp173 and Gly174 of EGFP in both orientations (Figure 1a). For the RRRWWW insertion, we deleted the two acidic residues in the loop, Glu172 and Asp173, which may otherwise partially neutralize the positive charges of the CPP and reduce its cell-penetrating activity. Fortuitously, in addition to the desired constructs, insertion mutagenesis also generated a construct containing an extra arginine residue, RRRRWWW, likely as the result of frame shift mutation during homologous recombination of the PCR products in bacterial cells. The EGFP insertion mutants generated in this study and their properties are summarized in Table 1. Both wild-type and mutant forms of EGFP were expressed in *E. coli* and purified to near homogeneity in high yields. Although the mutant proteins showed slightly reduced fluorescence intensity (10-50%) relative to wild type EGFP, their excitation and emission maxima remained essentially unchanged (Figure S1).

**Figure 1.**
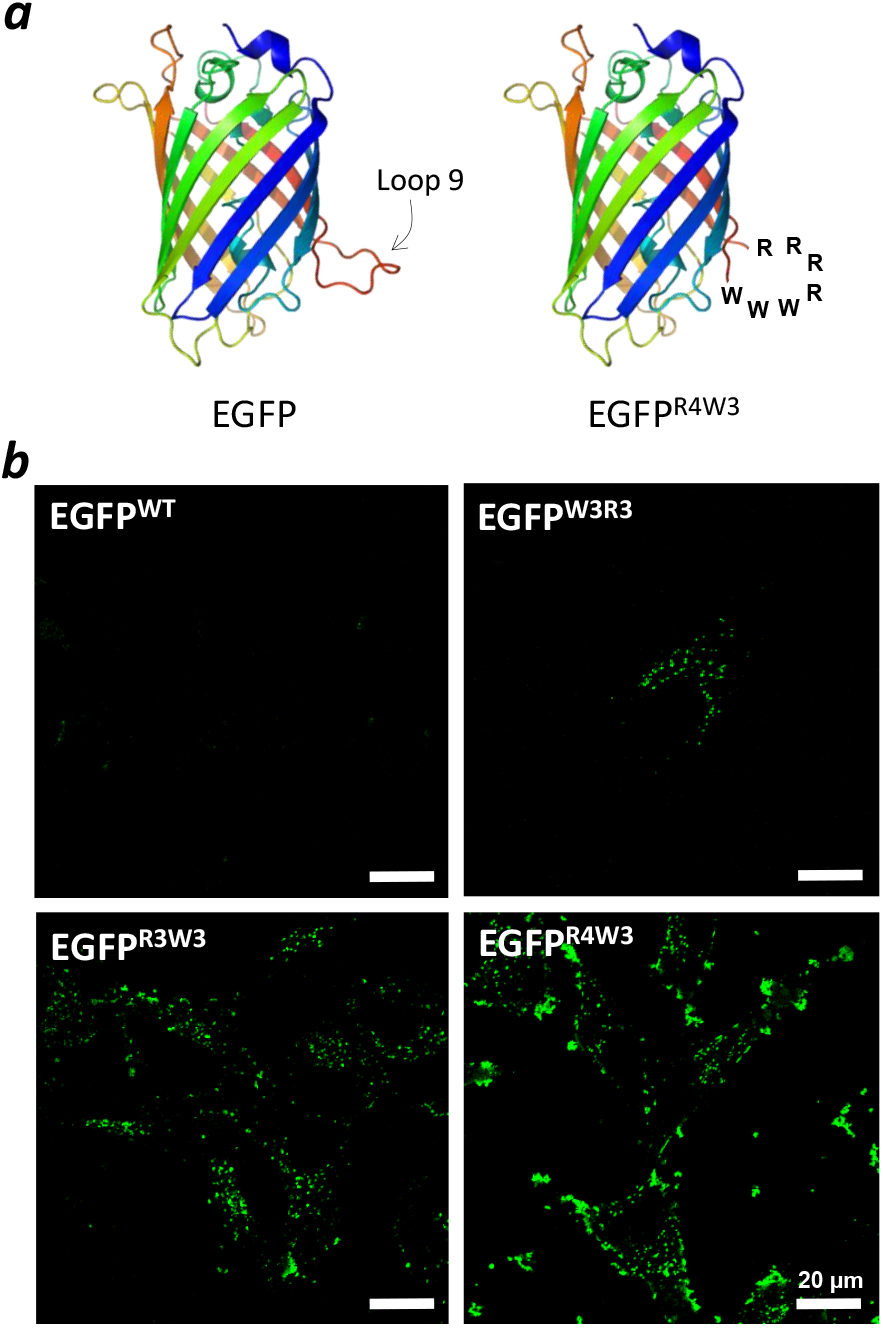
(a) Structures of WT and mutant EGFP showing the location of loop 9 and the inserted CPP motif. (b) Livecell confocal microscopy images of HeLa cells after treatment with WT or mutant EGFP (5 μM) for 2 h in the presence of 1% FBS.

**Table 1.**
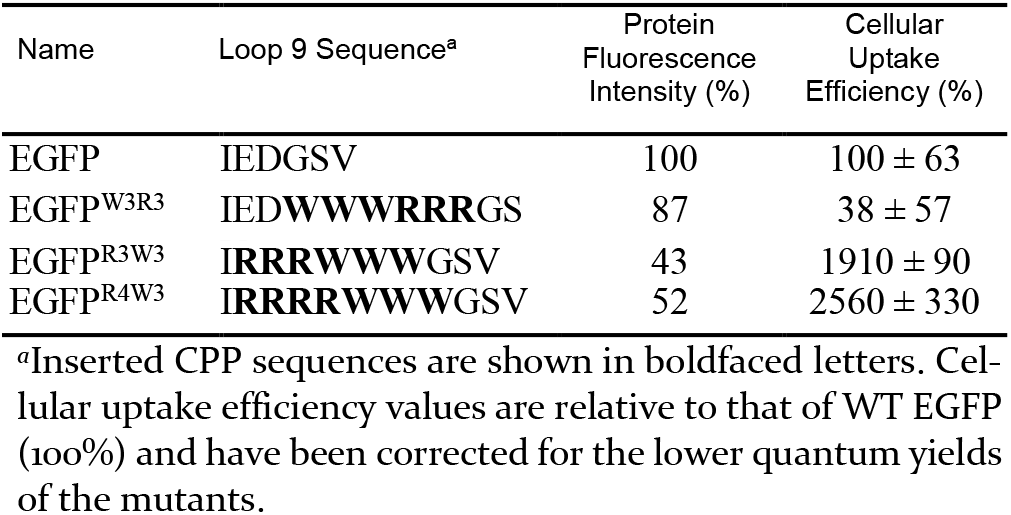
Structures and Properties of EGFP Variants ^a^

To determine the cellular entry efficiency of EGFP and the insertion mutants, HeLa cells were treated with 5 μM protein for 2 h in the presence of 10% fetal bovine serum (FBS), washed, and analyzed by flow cytometry. While EGFP^W3R3^ showed no significant improvement in cellular uptake compared to WT EGFP, EGFP^R3W3^ and EGFP^R4W3^ entered the cells with 19- and 26-fold higher efficiency than EGFP^WT^ (Table 1). To confirm the flow cytometry results, we also treated HeLa cells with the EGFP variants (5 μM) for 2 h and imaged the cells by live-cell confocal microscopy. The strongest fluorescence was observed in cells treated with EGFP^R4W3^, followed by EGFP^R3W3^ and EGFP^W3R3^, whereas cells treated with WT EGFP showed no detectable intracellular fluorescence (Figure 1b). The punctate fluorescence patterns observed for the three mutant proteins suggest that they were taken up by the cell through endocytosis and a substantial fraction of the internalized proteins remained entrapped inside the endosomes and lysosomes. Unfortunately, neither flow cytometry nor confocal microscopy revealed whether any of the internalized proteins reached the cytosol. The poorer cellular uptake of EGFP^W3R3^ than EGFP^R3W3^ is likely caused by the presence of two negatively charged residues adjacent to the CPP motif in the former (Table 1). It is also possible that the WWWRRR motif (when inserted into loop 9 of EGFP) binds the plasma membrane less effectively than RRRWWW.

### Generation of Cell-Permeable Protein Tyrosine Phosphatase 1B

To demonstrate cytosolic entry as well as the generality of our approach, we next chose to deliver the catalytic domain (amino acids 1-321) of protein-tyrosine phosphatase 1B (PTP1B) into mammalian cells. Tyrosine phosphorylation is generally restricted to cytosolic and nuclear proteins or the cytosolic domains of transmembrane proteins. Any perturbation of the phosphotyrosine (pY) levels of cellular proteins would therefore provide definitive evidence for functional delivery of PTP1B into the cytosolic space. Moreover, any change in the pY level can be conveniently detected by immunoblotting with an anti-pY antibody. Inspection of the PTP1B(1-321) structure^45^ revealed five solvent exposed loop regions as potential sites for CPP grafting (Table 2). These loops are distal from the catalytic or allosteric site of PTP1B. Sequence alignment with other members of the PTP family revealed a high degree of sequence variation in these loop regions,^46^ suggesting that modification of these loops is likely to be tolerated with regard to the folding and catalytic function of PTP1B. For each loop, the CPP sequence was inserted in both orientations, WWWRRRR and RRRRWWW, resulting in a total of 10 loop insertion mutants (Table 2). The mutant proteins were designated as “1-5W” and “1-5R”, based on the site of insertion (i.e., “1-5” for loops 1-5, respectively) and the CPP orientation (“W” for WWWRRRR and “R” for RRRRWWW). To ensure an overall positive charge at the modified loops, some of the acidic residues in the original loop regions were deleted. In some cases, glycine residues were added to both sides of the CPP sequence to maintain a minimal level of loop flexibility. The 3D structures of the 10 PTP1B mutants were predicted by using the online protein fold recognition server Phyre 2.^47^ All 10 mutants were predicted to have wild-type protein fold with the CPP sequences displayed at the protein surface (Figure S2). For loop 1, 3, and 5 insertion mutants, the CPP motifs adopted “cyclic-like” topology with the side chains facing the solvent, whereas in Loop 2 and 4 mutants, the CPPs showed a less constrained structure.

**Table 2.**
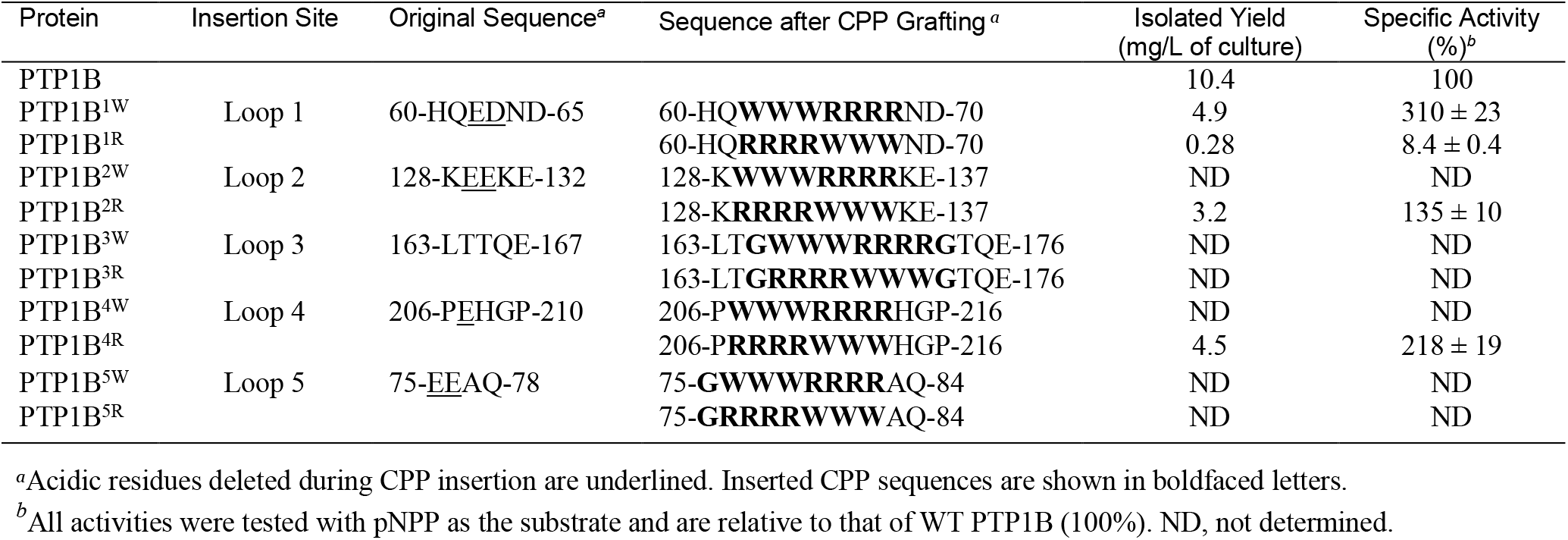
Structures and Properties of WT and Mutant PTP1B

The PTP1B mutants were generated by the one-step polymerase chain reaction method.^48^ To quickly assess their solubility and catalytic activity, each of the mutants was expressed in 5 mL of *Escherichia coli* cell culture and the crude cell lysates were analyzed by SDS-PAGE. When expressed at reduced temperature, all 10 insertion mutants produced predominantly soluble proteins indicating that insertion of CPP into the loops did not disrupt the global folding of PTP1B (Figure S3). The phosphatase activities in the cell lysates were quantitated by using *p*-nitrophenyl phosphate (pNPP; 0.5 mM) as the substrate. Four out of the 10 mutants (PTP1B^1W^, PTP1B^1R^, PTP1B^2R^, and PTP1B^4R^) showed catalytic activities that are 25-60% of wild type PTP1B, while the rest were much less active (Figure 2). Note that the total PTP activity in a cell lysate is governed by both the expression level and the specific activity of a given mutant.

**Figure 2.**
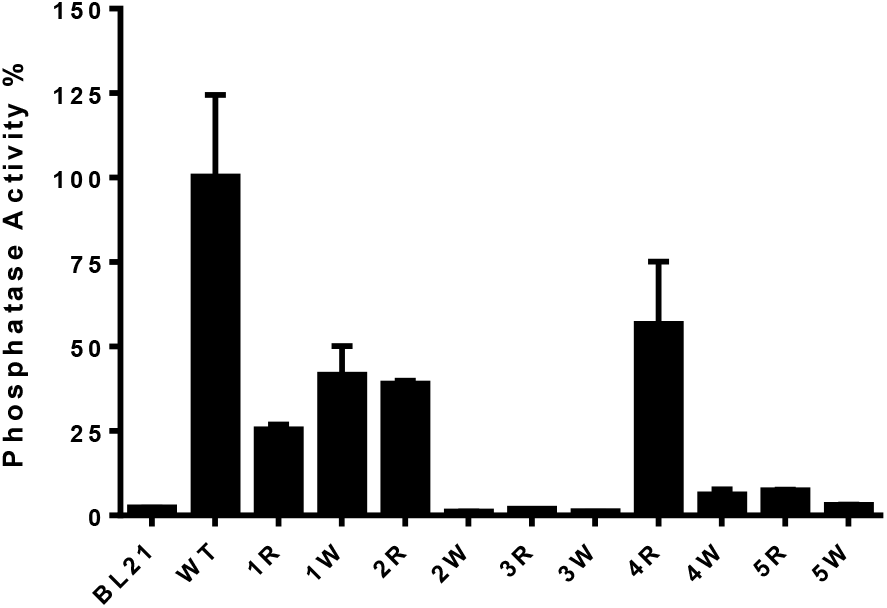
Phosphatase activity in the crude lysates of *E. coli* BL21(DE3) cells expressing WT or mutant PTP1B. Data shown represent the mean and SEM of three independent experiments and are normalized to that of WT PTP1B (100%). BL21, untransformed cells.

The four most active PTP1B mutants (PTP1B^1W^, PTP1B^1R^, PTP1B^2R^, and PTP1B^4R^) were expressed in *E. coli* cells on a large scale and purified to near homogeneity by metal affinity chromatography (all mutants contained an C-terminal histidine tag). The four mutants showed different yields of soluble protein, likely caused by different folding efficiency and proteolytic stabilities. The specific activities of the mutants were determined with the purified proteins and compared to that of wild type PTP1B. With the exception of PTP1B^1R^, which had both low yield and poor specific activity, the other three mutants were isolated in good yields and showed similar or higher catalytic activities to/than wild type PTP1B (Table 2).

To assess the cell permeability of the PTP1B mutants, NIH 3T3 fibroblast cells were treated with wild-type or mutant PTP1B for 2 h and lysed. The crude cell lysates were separated by SDS-PAGE and the gel was immunoblotted with anti-pY antibody 4G10. While the untreated cells and cells treated with wild-type PTP1B showed very similar pY protein patterns, the cells after treatment with PTP1B^2R^ and PTP1B^4R^ exhibited greatly reduced global pY levels (Figure 3A). Reduced pY levels were also apparent for cells treated with PTP1B^1W^ and PTP1B^1R^, although to a less extent compared to those of PTP1B^2R^ and PTP1B^4R^. Further, 3T3 cells treated with different concentrations of PTP1B^2R^ exhibited dose-dependent reduction of the pY level for most proteins, with some of the proteins (e.g., those at ~30, 37, 41, 47, and 55 kD) being more sensitive to PTP1B than others (Figure 3B). Similar effect on pY levels was also observed in HeLa cervical cancer and A549 lung cancer cells (Figure S4). These data indicate that the PTP1B mutants effectively entered the cytosol of mammalian cells and were biologically active in dephosphorylating tyrosine residues of intracellular proteins. These PTP1B variants may provide a useful tool for cell signaling research by globally reducing the pY levels of intracellular proteins.

**Figure 3.**
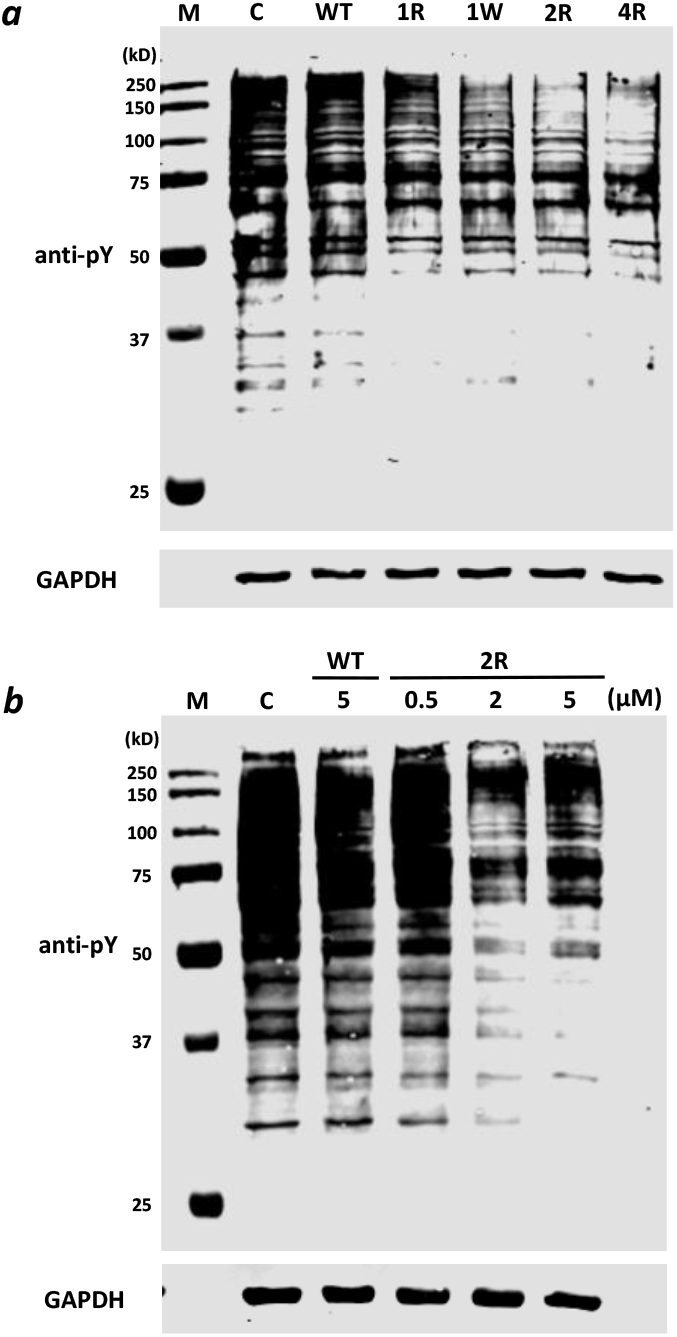
Effect of WT and mutant PTP1B on the global pY levels in NIH 3T3 cells. (***a***) Anti-pY Western blot analysis of NIH 3T3 cells after treatment for 2 h with wild-type or mutant PTP1B (2.1 μM for PTP1B^1R^ due to limited availability and 3.0 μM for all other proteins) in the presence of 1% serum. (***b***) Dose-dependent reduction of global pY levels as a function of PTP1B^2R^ concentration (0.5-5 μM). The membrane was re-blotted with anti-GAPDH antibody to ensure equal sample loading. M, molecular weight markers; C, control cells without PTP1B treatment.

### Intracellular Delivery of Purine Nucleoside Phosphorylase as Potential Enzyme Replacement Therapy

Purine nucleoside phosphorylase (PNP) is an enzyme involved in purine metabolism, by converting inosine into hypoxanthine and guanosine into guanine, plus ribose phosphate.^49^ Mutations that result in PNP deficiency cause defective T-cell (cell-mediated) immunity but can also affect B-cell immunity and antibody responses.^50^ A potential treatment of this rare genetic disease is to deliver enzymatically active PNP into the cytosol of patient cells. Examination of the homotrimeric structure of PNP^51^ revealed three solvent exposed loops that are also distal from the active site, namely His^20^-Pro^25^, Asn^74^-Gly^75^, and Gly^182^-Leu^187^. We inserted the CPP motif RRRRWWW into each of these loop regions to produce three PNP variants (Table 3). For the third insertion mutant (182-187), an acidic residue (Glu183) was removed to maximize overall positive charges at the loop sequence. Pilot expression experiments under different induction conditions revealed that CPP insertion at site 1 or 2 resulted in insoluble proteins, whereas insertion at site 3 produced a partially soluble protein, PNP^3R^, which was purified to near homogeneity following the same procedure as for wild-type PNP. PNP^3R^ has similar catalytic activity to the wild-type enzyme (Table 3).

**Table 3.**
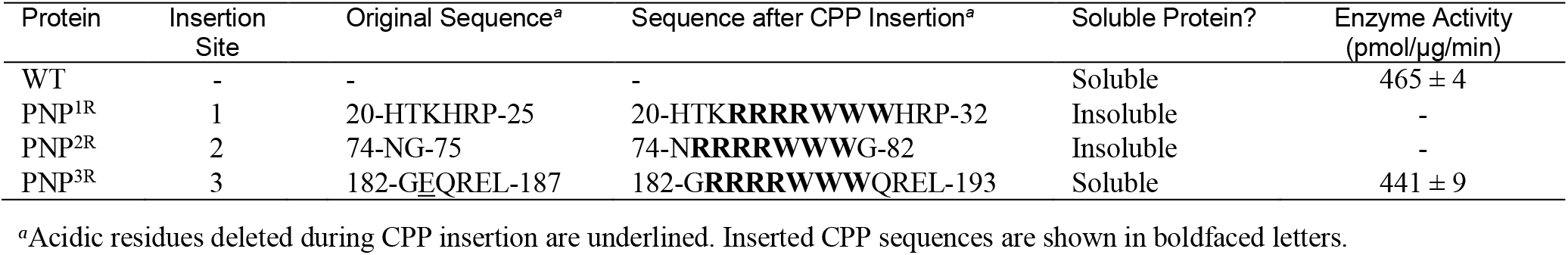
Structures and Properties of PNP Insertion Mutants

Cellular entry of PNP^3R^ was first examined by treating HeLa cells for 5 h with 5 μM fluorescein-labeled PNP^3R^ or wild-type PNP (PNP^WT^) and imaging the cells by live-cell confocal microscopy. Cells treated with PNP^3R^ showed readily visible green fluorescence signals inside the cells, whereas cells treated with PNP^WT^ showed no detectable fluorescence under the same experimental condition (Figure 4a). Note that the proteins were intentionally labeled at a low stoichiometry (0.1-0.2 dye/protein) to minimize any protein precipitation or denaturation. To further assess the cellular entry efficiency of PNP^3R^, PNP-deficient mouse T lymphocytes (NSU-1) were treated with 1 μM PNP^WT^ or PNP^3R^ for 2 h and washed exhaustively to remove extracellular proteins. The cells were lysed and the PNP activities in cytosolic fractions were quantified by using a commercial PNP enzymatic assay kit. While the untreated NSU-1 cells had no significant PNP activity, treatment of NSU-1 cells with PNP^3R^ resulted in 1.35-fold higher PNP activity than that of normal S49 cells (100%; Figure 4b). Under the same condition, NSU-1 cells treated with PNP^WT^ showed an activity that was 16% relative to that of S49 cells. The latter activity is likely due to incomplete removal of the extracellular PNP activity by the washing procedure, as NSU-1 cells are non-adherent cells and it was difficult to remove the extracellular fluids completely during washing.

**Figure 4.**
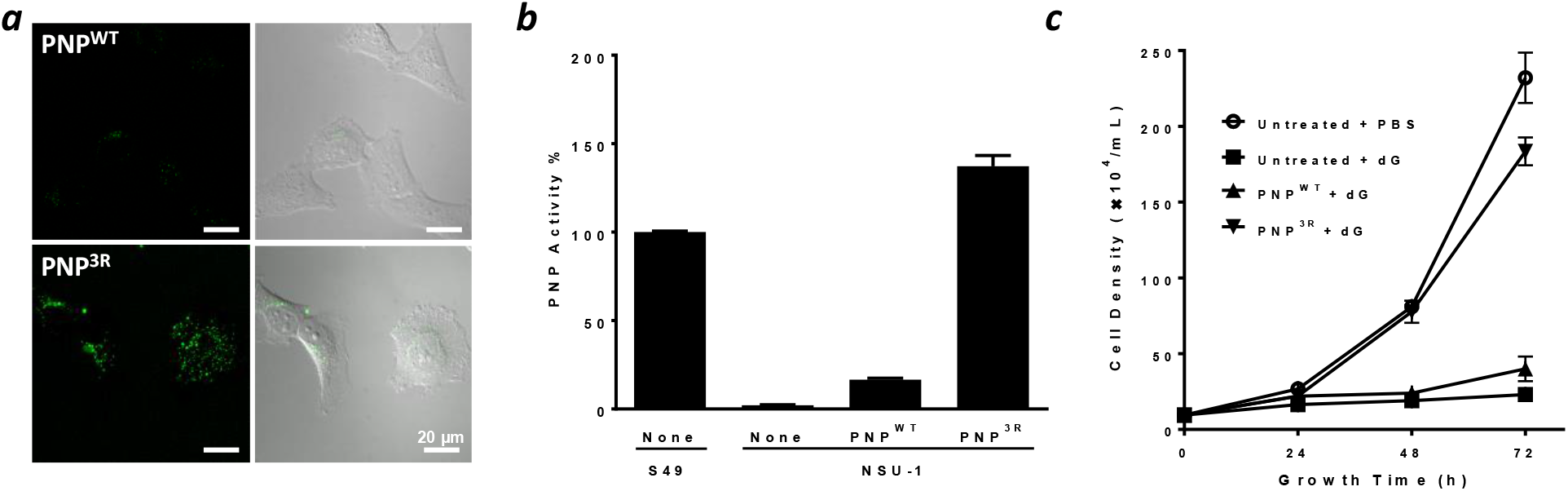
Cellular entry and biological activity of PNP^3R^. (***a***) Live-cell confocal images of HeLa cells after treatment with 5 μM fluorescein-labeled PNP^WT^ (top) or PNP^3R^ (bottom) for 5 h in the presence of 1% FBS. Left panels, FITC fluorescence; right panels, overlap of FITC signals with the DIC images of the same cells. (***b***) PNP activities in cell lysates derived from S49 (wildtype PNP) or NSU-1 cells with and without treatment with PNP^WT^ or PNP^3R^ (1 μM). Representative data (mean ± SD) from three independent experiments are shown. (***c***) Protection of NSU-1 cells against dG toxicity. NSU-1 cells were treated with PBS (no protein), 3 μM PNP^WT^, or 3 μM PNP^3R^ for 6 h at 37 °C, washed exhaustively, and incubated with trypsin-EDTA for 3 min. The cells were seeded at a density of 1 x 10^5^ cells/mL in DMEM containing 25 μM dG and cell growth (cell counts) was monitored for 72 h. Cells without protein or dG treatment were used as positive control.

Finally, we tested the capacity of PNP^3R^ to correct the metabolic defects of NSU-1 cells caused by PNP deficiency. PNP-deficient cells (e.g., NSU-1) are sensitive to deoxyguanosine (dG) toxicity. As shown in Figure 4c, NSU-1 cells failed to grow in the presence of 25 μM dG, while in the absence of dG the cell density increased from 1 x 10^5^ to 2.3 x 10^6^ cells/mL in 72 h. When NSU-1 cells were pretreated with 3 μM PNP^3R^ for 6 h, washed exhaustively to remove any extracellular PNP^3R^, and then challenged with 25 μM dG, they exhibited a growth curve similar to that of the untreated cells (no dG, no protein). Under the same conditions, NSU-1 cells treated with PNP^WT^ showed only a small amount of growth (13%) relative to the untreated control, likely due to incomplete removal of PNP^WT^ from the growth medium. Thus, PNP^3R^, but not PNP^WT^, can effectively rescue PNP-deficient cells against dG toxicity. PNP^3R^ may be further developed into a novel, intracellular enzyme replacement therapy. All previous enzyme replacement therapies involved extracellular or lysosomal enzymes.^52^

### Serum Stability of Loop Insertion Mutants

Insertion of amphipathic CPP sequences (e.g., RRRRWWW) into surface loops may decrease the thermodynamic stability of a protein as well as generates potential new cleavage sites for proteases (e.g., trypsin and chymotrypsin). Both factors can potentially reduce the metabolic stability of the mutant proteins. We thus tested the proteolytic stabilities of wild-type EGFP, PTP1B, and PNP as well as the biologically active mutants derived from this study by incubating them in human serum for varying periods of time (0-16 h) and quantitating the amounts of remaining intact protein by SDS-PAGE analysis (Figure S5). The wild-type proteins were all highly stable in serum, exhibiting *t*_1/2_ values of >16 h (Figure 5). Among the seven mutant proteins tested, six showed comparable or slightly reduced stability relative to their wild-type counterparts; only one mutant, PTP1B^1W^, showed substantially more rapid degradation than wild-type PTP1B. We also examined the serum stability of PNP^WT^ and PNP^3R^ by monitoring their enzymatic activities as a function of the incubation time and observed no significant loss of PNP activity for either protein after 16 h of incubation (Figure S6). Since linear CPPs (e.g., Phe-Nal-Arg-Arg-Arg-Arg) are highly susceptible to proteolytic degradation (serum *t*_1/2_ ≤30 min),^33,53^ these data demonstrate that insertion of short CPP sequences into protein loops greatly increases their proteolytic stability, likely as a result of restrained conformations.

**Figure 5.**
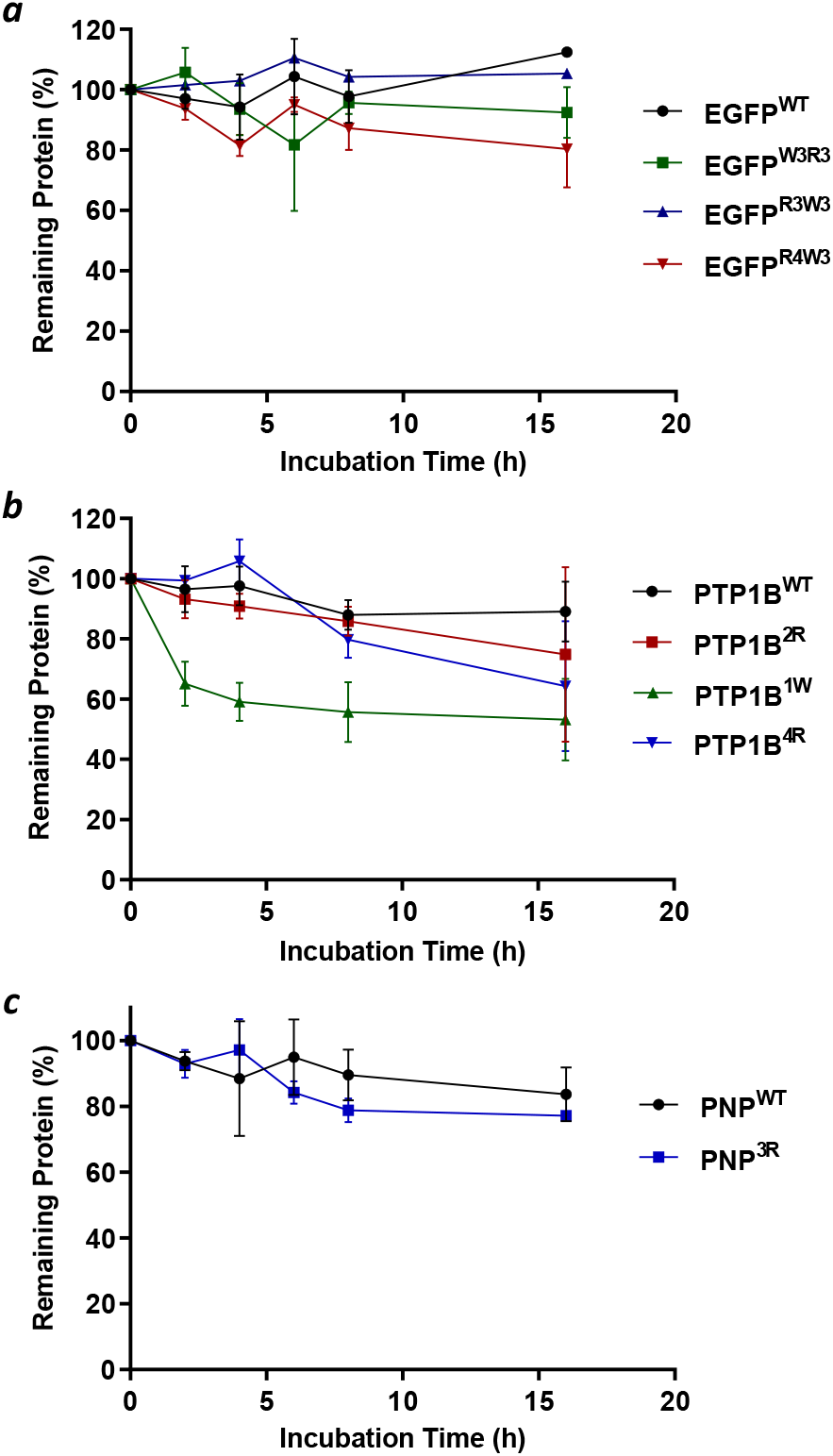
Serum stability of wild-type and mutant forms of EGFP (*a*), PTP1B (*b*), and PNP (*c*). Values reported represent the amounts of intact protein remaining at indicated time points and are relative to that at time zero (100%).

### Comparison to Other Protein Delivery Methods and Mechanistic Implications

The loop-insertion approach offers a number of advantages over other reported protein engineering/delivery methods, not the least of which is its simplicity. In our case, once a recombinant protein is purified (e.g., from an *E. coli* cell lysate), it may be used directly as a biological probe or therapeutic agent. Additionally, whereas post-translational conjugation of a protein with a CPP (or other chemical entities) often results in a mixture of different species, the loop-insertion method produces a single species of well-defined structure. Compared to other protein surface remodeling methods (e.g., supercharging^19,20^ and esterification^21^), our method involves relatively minor changes to the protein structure and should be applicable to a broader range of proteins. The resulting mutant proteins are more likely to retain the original protein fold/activity and less likely to elicit immune responses. Finally, the CPP motifs grafted to protein loops are structurally constrained and relatively stable against proteolytic degradation, whereas fusion of CPP sequences to the N- or C-terminus of a protein, as practiced by previous investigators, generates recombinant proteins that frequently lack sufficient metabolic stability for clinical applications.

Compared to posttranslational conjugation of proteins with some of the cyclic CPP(s),^33,35^ the loop-insertion mutants from this work appear to have lower cytosolic entry efficiencies. When examined by confocal microscopy, the cell-permeable proteins derived from this study produced punctate intracellular fluorescence patterns (Figures 1b and 4a), indicating incomplete endosomal escape of these proteins. We envision that their endocytic uptake and/or endosomal escape efficiency may be improved by exploring other proteinogenic amino acids (in addition to Arg and Trp) for the CPP motif as well as different insertion sites on the protein surfaces. After all, bacteria and viruses have evolved highly efficient systems to deliver their proteins and nucleic acids into the cytosol of mammalian cells. Non-proteinogenic amino acids (e.g., Nal) may also be leveraged to improve the efficiency of the CPP motifs.^42^ We have previously shown that substitution of Nal for Phe (or Trp) greatly improves the cytosolic entry efficiency of cyclic CPPs [e.g., cyclo(Phe-Nal-Arg-Arg-Arg-Arg-Gln) and cyclo(Phe-phe-Nal-Arg-arg-Arg-arg-Gln)].^32,34^

Our study also has important mechanistic implications. As discussed above, proteins derived from diverse organisms (from bacteria to man) have demonstrated the capacity to enter the cytosol of mammalian cells, but their mechanism(s) of cellular entry has not yet been resolved. In the case of bacterial toxins, it is well established that the proteins are brought into the early endosome by receptor-mediated endocytosis and some of the toxins are capable of directly crossing the endosomal membrane into the cytosol.^54,55^ The mechanism of endosomal escape, however, remains controversial despite decades of intensive studies. A popular hypothesis states that, as the endosomal pH decreases, the transmembrane domain of the toxin undergoes a conformational change and inserts into the endosomal membrane to form an α-helical or β-barrel pore, through which the effector domain translocates from the endosome to the cytosol in its denatured state.^54,55^ Although our current findings do not disprove the pore-formation mechanism (for bacterial toxins) per se, they demonstrate that proteins are capable of escaping the endosome by a mechanism(s) other than pore formation. None of the proteins examined in this work (EGFP, PTP1B, and PNP) are known to undergo acid denaturation and membrane insertion to form a pore. In fact, PTP1B is very stable at the endosomal pH (4.5-6.5) and catalytically most active at pH 5.5.^56^ It is also highly unlikely that insertion of RRRRWWW into surface loops would endow the above proteins the capability of pore formation. Rather, we believe that the inserted CPP motif functions as a “cyclic CPP” – by binding first to the plasma membrane to facilitate endocytic uptake of the proteins and then to the endosomal membrane to induce vesicle budding and collapse.^41^ We further hypothesize that, like the cyclic CPPs, bacterial toxins may exit the endosome by inducing vesicle budding and collapse from the endosomal membrane.^57^

## CONCLUSION

In this study, we have demonstrated that proteins may be rendered cell-permeable by inserting a short CPP motif into one of their surface loop regions. For the three proteins investigated in this study (EGFP, PTP1B, and PNP), a total of 16 mutants were generated (Table 4). Among these mutants, eight (50%) produced soluble proteins in good yields, seven of which (44%) exhibited wild type-like biochemical activity and six of which (38%) were cell-permeable. Importantly, for each of the three proteins tested in this work, at least one soluble, cell-permeable, and biologically active mutant was obtained. We conclude that insertion of CPP motifs into the surface loops of proteins represents a general approach to engineering cell-permeable proteins.

**Table 4.** Summary of Loop Insertion Mutants Generated in This Study

| Protein | No. of Mutant Constructs Generated | No. of Soluble Proteins | No. of Proteins with WT-Like Activity | No. of Cell-Permeable Proteins |
| --- | --- | --- | --- | --- |
| EGFP | 3 | 3 | 3 | 2 |
| PTP1B | 10 | 4 | 3 | 3 |
| PNP | 3 | 1 | 1 | 1 |
| Total | 16 | 8 | 7 | 6 |

## ASSOCIATED CONTENT

### Supporting Information

Experimental details and additional data are available free of charge online at http://pubs.acs.org.

## AUTHOR INFORMATION

### Author Contributions

D.P. conceived the project, D.P. and K.C. designed the experiments, K.C. carried out the experiments, and D.P. and K.C. wrote the manuscript.

### Notes

The authors declare no competing financial interests.

## ACKNOWLEDGMENT

We thank Dr. Z. Qian and A. Sahni for their technical assistance during confocal microscopy experiments.

## ABBREVIATIONS

CPP: cell-penetrating peptide
dG: deoxyguanosine
EGFP: enhanced green fluorescent protein
PNP: purine nucleoside phosphorylase
PTP: protein tyrosine phosphatase.

---

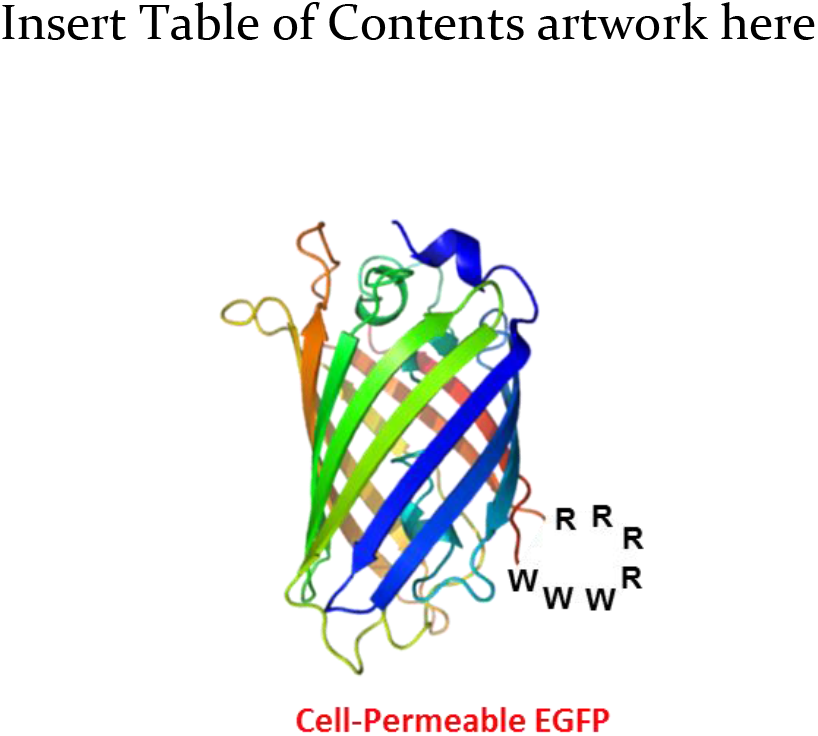

