## Supplementary Materials for "Engineering Cell-Permeable Proteins through Insertion of Cell-Penetrating Motifs into Surface Loops"

### Supporting Information

### Table of Contents

**Experimental Procedures** .......... S1
**Table S1** .......... S7
**Table S2** .......... S8
**Figure S1** .......... S10
**Figure S2** .......... S10
**Figure S3** .......... S11
**Figure S4** .......... S11
**Figure S5** .......... S12
**Figure S6** .......... S12
**Figure S7** .......... S13

### EXPERIMENTAL PROCEDURES

**Materials.** Plasmid encoding histidine-tagged human PNP was a gift from Dr. Vern Schramm of Albert Einstein College of Medicine (Addgene plasmid # 64076).[1] Murine T lymphoma NSU-1 cell line was a gift from Dr. R. Scott McIvor, University of Minnesota. Cell culture media, fetal bovine serum (FBS), penicillin-streptomycin, 0.25% trypsin-EDTA, DPBS (2.7 mM potassium chloride, 1.5 mM monopotassium phosphate, 8.9 mM disodium hydrogen phosphate, and 137 mM sodium chloride) were purchased from Invitrogen (Carlsbad, CA). Fluorescein isothiocyanate (FITC), 5(6)-carboxynaphthofluorescein succinimidyl ester (NF-NHS ester), 2'-deoxyguanosine, imidazole, protease inhibitor cocktail, ampicillin, lysozyme, oligonucleotides, tryptone enzymatic digest, and yeast extracts were purchased from Sigma-Aldrich (St. Louis, MO). Slide-A-Lyzer Dialysis Cassettes were purchased from Thermo Fisher Scientific (Waltham, MA). Restriction enzymes and DNA polymerase were purchased from New England BioLabs (Ipswich, MA). Purine nucleoside phosphorylase activity assay kit (Fluorogenic) was purchased from Abcam (Cambridge, MA). HisTrap FF nickel affinity column was purchased from GE Healthcare (Marlborough, MA). Protein assay dye (Bradford reagent) and Micro Bio-Spin 6 desalting columns were purchased from Bio-Rad (Hercules, CA). Isopropyl β-D-1-thiogalactopyranoside (IPTG) was purchased from Teknova Inc (Hollister, CA). Ampicillin was purchased from Cayman Chemical Company (Ann Arbor, MI). Tris(2-carboxyethyl)phosphine hydrochloride (TCEP) and bovine serum albumin (BSA) were purchased from VWR (West Chester, PA). All solvents and other chemical reagents were obtained from Sigma-Aldrich, Fisher Scientific (Pittsburgh, PA), or VWR (West Chester, PA) and were used without further purification unless noted otherwise.

**Mutagenesis.** Insertion of CPP sequences into loop regions of target proteins was carried out by following a modified one-step overlapping extension PCR protocol.[2] For each mutant, a pair of DNA primers was designed, with each primer containing a 5' overlapping region that encodes the desired CPP sequence and a 3' annealing region that is complementary to the DNA sequence before or after the insertion site (**Table S1**). Deletion of native loop residues was effected by removing their respective codons from the primer annealing region. The DNA primers were purchased from Sigma and used to amplify the plasmid DNA encoding the target protein by polymerase chain reaction. The PCR product was treated with restriction enzyme *Dpn*I and used to transform *E. coli* DH5α competent cells. The linear PCR product was cyclized in *E. coli* via homologous recombination. Positive clones harboring the intended insertions were identified by restriction mapping and the DNA sequences of the entire protein coding regions were confirmed by Sanger sequencing. The amino acid sequences of the wild-type and mutant proteins used in this work are listed in **Table S2**.

**Expression and Purification of EGFP Mutants.** Wild-type and mutant EGFP containing a C-terminal six-histidine tag were expressed in *Escherichia coli* BL21(DE3) cells. *E. coli* cells harboring the proper plasmid were grown in Luria Broth supplemented with 75 μg/mL ampicillin at 37 °C until $OD_{600}$ reached 0.6, when protein expression was induced by the addition of 0.2 mM IPTG at 16 °C for 20 h. The cells were pelleted by centrifugation and suspended in 50 mL (for 1 L of cell culture) of Tris buffer (50 mM Tris, pH 8.0, 300 mM NaCl, 3 mM beta-mercaptoethanol) supplemented with 0.2 mg/mL lysozyme and protease inhibitor cocktail (Sigma). After incubation at 4 °C for 30 min, the cell lysate was briefly sonicated and clarified by centrifugation at 20,000 g for 45 min. The cell lysate was loaded onto a "HisTrap FF 5mL" nickel affinity column (GE Healthcare) on an AKTA explorer FPLC system (Amersham Pharmacia Biotech). After washing with 50 mM imidazole, bound protein was eluted with a linear gradient of 50–500 mM imidazole in the Tris buffer described above.

For wild-type EGFP, elution fractions were combined, concentrated in centrifugal filter units (Millipore), and dialyzed against PBS. For EGFP mutants, the elution fractions were combined and the purified proteins were precipitated by the slow addition of ammonium sulfate to 50% saturation. Protein precipitates were re-dissolved in a minimal volume of the Tris buffer and dialyzed against PBS. Protein purity was assessed by SDS-PAGE and judged to be ≥95% (**Figure S7a**). Protein concentration (typically 1.5 - 12 mg/mL) was determined by the Bradford assay. The purified protein was supplemented with 20% glycerol, aliquoted, quickly frozen and stored at -80 °C.

**Protein Folding Prediction.** Protein folding prediction of PTP1B mutants were generated by online server Phyre2 (**P**rotein **H**omology/analog**Y** **R**ecognition **E**ngine V 2.0).[3] Folding prediction was carried out under normal modeling mode. Wild-type PTP1B catalytic domain (1-321) structure was chosen as the highest scoring template for all loop insertion mutants. Protein structures were analyzed and depicted by PyMOL (**Figure S2**).

**Quantitation of PTP1B Activity in Crude *E. coli* Lysates.** *E. coli* BL21(DE3) cells were transformed with the proper PTP1B gene and plated on Luria Broth agar. For each mutant, three different colonies were randomly picked and each was used to inoculate 5 mL of Luria Broth supplemented with 75 µg/mL ampicillin. Cells were grown at 37 °C until $OD_{600}$ reached 0.6, when protein expression was induced by the addition of 0.2 mM IPTG (final concentration) at 18 °C for 20 h. Cells were harvested by centrifugation at 3500 rpm for 10 min, and suspended in 0.5 mL of lysis buffer [40 mM HEPES (pH 7.5), 300 mM NaCl, 1 mM TCEP] supplemented with 0.2 mg/mL lysozyme, 2 mM PMSF, and protease inhibitor cocktail (Roche). After incubation for 30 min at 4 °C, the mixture was sonicated (20 pulses, 1 second per pulse) and centrifuged at 20,000 g for 10 min. The clear supernatant was removed by a micropipette and transferred into a clean Eppendorf tube, while the insoluble fraction was re-suspended in an equal volume (0.5 mL) of the lysis buffer. Both soluble and insoluble fractions of the cell lysate were analyzed by SDA-PAGE to check the expression level and solubility. To assess the phosphatase activity in the cell lysate, 5 µL of the soluble fraction was added into 150 µL of reaction mixture containing 50 mM HEPES (pH 7.0), 50 mM NaCl, 1 mM EDTA, 2 mM TCEP, and 0.5 mM *p*-nitrophenyl phosphate (pNPP). The reaction was incubated in a 96-well plate at 37 °C for 20 min, before being quenched by the addition of 150 µL of 1 M NaOH. Product formation was quantified by measuring the absorbance at 405 nm and all activities were normalized to that of cells expressing wild-type PTP1B. Untransformed BL21(DE3) cells were used as the negative control.

**Expression and Purification of PTP1B Mutants.** *E. coli* BL21(DE3) cells transformed with wild-type or mutant PTP1B DNA (with a C-terminal six-histidine tag) were grown in Luria Broth supplemented with 75 µg/mL ampicillin at 37 °C until $OD_{600}$ reached 0.6. Protein expression was induced by the addition of 0.2 mM IPTG at 18 °C for 20 h. The cells were pelleted by centrifugation at 3500 rpm for 15 min. The cell pellets (from 1 L culture) were suspended in 50 mL of lysis buffer (40 mM HEPES, pH 7.5, 300 mM NaCl, 3 mM beta-mercaptoethanol), supplemented with 0.2 mg/mL lysozyme, 2 mM PMSF and protease inhibitor cocktail (Roche). After incubation at 4 °C for 30 min, the cell lysate was sonicated and clarified by centrifugation at 20,000 g for 45 min. The clear cell lysate was loaded onto a "HisTrap FF 5 mL" nickel affinity column (GE Healthcare) on an AKTA explorer FPLC system (Amersham Pharmacia Biotech). After washing with 50 mM imidazole, bound protein was eluted with the lysis buffer containing a linear gradient of 50-500 mM imidazole (pH 8.0). Elution fractions containing pure PTP1B were combined and concentrated in centrifugal filter units (Millipore), followed by overnight dialysis against the lysis buffer to remove imidazole. Any precipitate formed during protein concentration and dialysis was removed by centrifugation. Protein purity was assessed by SDS-PAGE and judged to be ≥90% (**Figure S7b**). Protein concentration was determined by the Bradford assay. Protein aliquots were quickly frozen and stored at -80 °C.

**Phosphatase Activity Assay of Purified PTP1B Mutants.** The specific activities of wild-type and mutant PTP1B were measured in 150-μL reactions containing 50 mM HEPES (pH 7.0), 50 mM NaCl, 1 mM EDTA, 1 mM TCEP and 0.5 mM *p*-nitrophenyl phosphate (pNPP). The reaction was carried out at room temperature in a quartz microcuvette and initiated by the addition of PTP1B (final 50 nM). Product formation was continuously monitored at 405 nm in a UV-VIS spectrophotometer. Initial rates were calculated from the early phases of the reaction progress curves (≤60 s) and normalized to that of wild-type PTP1B.

**Expression and Purification of Wild-Type and Mutant PNP.** *E. coli* BL21(DE3) cells harboring the proper PNP expression vector containing an N-terminal six-histidine tag were grown in Luria Broth supplemented with 75 μg/mL ampicillin at 37 °C until $OD_{600}$ reached 0.6. For wild-type PNP, protein expression was induced with 0.1% lactose at 37 °C for 9 h. For $PNP^{3R}$, protein expression was induced with 0.1% lactose at 20 °C for 16 h. The cells (from 1 L culture) were pelleted and suspended in 50 mL lysis buffer (PBS containing 300 mM NaCl, pH 8.0, 10% glycerol, 3 mM beta-mercaptoethanol), supplemented with 0.2 mg/mL lysozyme, 2 mM PMSF and a protease inhibitor cocktail (Roche). After incubation at 4 °C for 30 min, the cell lysate was sonicated and clarified by centrifugation at 20,000 g for 45 min. The clear cell lysate was loaded onto a "HisTrap FF 5mL" nickel affinity column (GE Healthcare) on an AKTA explorer FPLC system (Amersham Pharmacia Biotech). After washing with 50 mM imidazole, bound protein was eluted with the lysis buffer containing a linear gradient of 50-500 mM imidazole (pH 8.0). Elution fractions containing pure PNP were combined and quickly frozen to -80 °C in aliquots. Protein purity was assessed by SDS-PAGE and judged to be ≥95% (**Figure S7c**). Protein aliquots were thawed and exchanged to the desired buffer by using Micro Bio-Spin 6 desalting column (Bio-Rad) before use. Enzyme activities of $PNP^{WT}$ and $PNP^{3R}$ were measured by using a fluorometric PNP activity assay kit (Abcam) following the manufacturer's instructions.

**Fluorescent Labeling of PNP.** The pH of protein solution in PBS (1–5 mg/mL) was adjusted to 8.5 with a $NaHCO_3$ solution. One equiv of an amine-reactive fluorescent dye (FITC) was slowly added and the reaction was allowed to proceed in the dark at at 4 °C overnight. The excess dye was removed by passing the reaction mixture through a desalting spin column (Bio-Rad). The dye/protein ratio was determined by comparing the absorbances at 280 nm and $\lambda_{max}$ of the FITC (495 nm).

**Cell Culture.** HeLa, NIH 3T3, S49, and NSU-1 cells were cultured in DMEM medium containing 10% FBS and 1% penicillin/streptomycin. All cell cultures were maintained in a humidified incubator at 37 °C in the presence of 5% $CO_2$.

**Flow Cytometry.** HeLa cells (1.5 x $10^5$ cells/mL) cultured in DMEM containing either 1% or 10% FBS in 12-well plates were treated with 5 μM wild-type or mutant EGFP at 37 °C for 2 h. Cells were washed twice with cold DPBS, detached from plate by trypsinization, and pelleted at 300 g for 5 min. The cells were washed two more times with cold DPBS, suspended in 200 μL of DPBS and analyzed on a BD FACS LSR II flow cytometer. For wild-type and mutant EGFP, the FITC channel was used with excitation wavelength of 488 nm. For naphthofluorescein (NF)-labeled peptides or proteins, the APC channel was used with excitation wavelength of 633 nm. Data presented were the mean ± SD of three independent experiments.

**Confocal Microscopy.** HeLa cells (5 x $10^4$ cells/mL) were seeded in a 35-mm glass-bottomed dish (Grenier Bio-One) and cultured overnight at 37 °C. The cells were washed twice with DPBS and treated with 5 μM EGFP or fluorescently labeled PNP in phenol red-free DMEM containing 1% FBS at 37 °C for 2-5 h. The cells were washed twice with DPBS, supplemented with phenol red-free DMEM, and imaged on a Nikon A1R live cell imaging confocal microscope. NIS Elements was used for image analysis and processing. Images presented in this report were generated using the instruments and services at the Campus Microscopy and Imaging Facility (Ohio State University).

**Immunoblotting.** NIH 3T3 cells ($10^6$ cells/well) were seeded in a 6-well plate one day before experiment. The cells were starved in serum-free medium for 3 h, followed by treatment with wild-type or mutant PTP1B (0-5 μM) for 2 h in medium containing 1% serum. Before harvest, 1 mM sodium pervanadate was added to the cells followed by incubation for 30 min. The cells were washed twice with cold DPBS, detached by trypsinization, and pelleted by centrifugation at 5,000 rpm for 10 min. Cells from each sample were then lysed on ice for 30 min in 100 μL of Pierce RIPA buffer (Thermo) supplemented with protease and phosphatase inhibitor cocktails (Thermo). The cell lysates were clarified by centrifugation at 20,000 g for 15 min, and the protein concentrations in the supernatants were determined by using BCA Protein Assay Kit (Thermo). Equal amounts of protein from each sample (30 μg) were separated by SDS-PAGE (10% polyacrylamide gel) and electrophoretically transferred to a nitrocellulose membrane at 100 V for 60 min. The membrane was blocked by 1-h incubation with 5% BSA in TBST, washed 3 times, and blotted with anti-pY monoclonal antibody 4G10 (1:1000 dilution, Millipore) at 4 °C overnight. After washing with TBST three times, the membrane was incubated with IRDye 800CW Goat anti-Mouse secondary antibody (1:10000 dilution, LI-COR) at room temperature for 2 h. The membrane was washed three times and scanned in the 800-nm channel for signal detection in an Odyssey CLx Imager (LI-COR). For GAPDH blotting, the same membrane was blocked again with 5% BSA in TBST at room temperature for 1 h, incubated with anti-GAPDH rabbit monoclonal antibody (1:5000 dilution, Cell Signaling Technology) at room temperature for 1 h, washed, and incubated with IRDye 800CW Goat anti-Rabbit secondary antibody (1:10000 dilution, LI-COR) at room temperature for 2 h, followed by fluorescence detection.

**Quantitation of Intracellular PNP$^{3R}$.** S49 and NSU-1 cells were seeded in a 96-well plate ($10^5$ cells/well) in 200 μL of medium. NSU-1 cells were treated with 1 μM PNP$^{WT}$ or PNP$^{3R}$ in DMEM containing 1% FBS at 37 ˚C for 2 h, while S49 cells remained untreated. The cells were pelleted at 300 g for 5 min and washed 5 times with 500 μL of cold DPBS. The cells were lysed in 40 μL of cytosolic lysis buffer (50 μg/mL digitonin, 75 mM NaCl, 1 mM NaH2PO4, 8 mM Na2HPO4, and 250 mM sucrose) on ice for 10 min, followed by centrifugation at 16,000 g for 5 min. The total protein concentrations in the cell lysates were determined by the BCA assay and adjusted to 20 μg/mL with a PNP assay buffer (Abcam). The PNP enzyme activities in the cell lysates were measured by using a fluorometric PNP activity assay kit (Abcam) and normalized to the PNP activity in S49 cell lysates.

**NSU-1 Growth Inhibition Assay.** NSU-1 cells (5 x $10^5$ cells) cultured in a 24-well plate were treated with 3 μM PNP$^{WT}$ or PNP$^{3R}$ in DMEM containing 1% FBS at 37 ˚C for 6 h. The cells were pelleted and washed twice with 1 mL of DPBS. Any extracellular protein was removed by incubation of cells with trypsin-EDTA (0.25 %, 2.21 mM) solution at 37 ˚C for 3 min, followed by additional washing with growth medium. The cells were then seeded in a 24-well plate at a starting density of 1 x $10^5$ cells/mL in DMEM containing 10% FBS, 1% penicillin/streptomycin, in the absence or presence of 25 μM 2'-deoxyguanosine. The cells were cultured at 37 ˚C for 72 h and the cell density was counted every 24 h.

**Serum Stability Assay**. Human serum was diluted to 50% in PBS and centrifuged at 15,000 rpm for 10 min to remove lipids. The clarified serum was mixed with a protein of interest in a total volume of 100 μL to give final concentrations of 25% serum and 10 μM protein. The mixture was incubated at 37 °C and 10-μL aliquots were withdrawn at different time points. The aliquots were immediately mixed with 10 μL of 2x SDS loading buffer, boiled for 5 min, and store at -20 °C. After incubation for up to 16 h, all aliquots were resolved on a 10% SDS-PAGE gel and the gel was stained with Coomassie Blue (**Figure S5**). The gel was scanned on an Odyssey CLx Imager (LI-COR) in the 700-nm channel and the band intensities (abundance) of interested protein was quantified by using ImageJ. All protein abundances were normalized to that of the sample at time zero. Assays were performed in triplicates and the mean ± SD values of protein abundance were plotted against the incubation time.

Alternatively, 40-μL reactions containing 25% human serum and 2 μM wild-type or mutant PNP protein in DPBS were set up 16, 8, 6, 4, 2 and 0 h before PNP activity assay and incubated at 37 °C. Five μL of each reaction mixture was withdrawn and diluted into 200 μL of PNP assay buffer (Abcam), and 50 μL of the diluted solution was used to measure any remaining PNP enzyme activity by using a fluorometric PNP activity assay kit (Abcam, final protein concentration = 25 nM). The remaining PNP activity was calculated from the early phase of the reaction progress curve (the initial linear range) and normalized to that of 0-h incubation. Assays were performed in triplicates and the mean ± SD values of enzyme activity were plotted against the incubation time (**Figure S6**).

**Table S1.** List of Primers Used for Constructing Loop Insertion Mutants

| **Protein Mutant** | **Oligonucleotide Sequences (5' to 3')** |
|---|---|
| EGFP$^{W3R3}$ | **TGGTGGTGGCGCCGCCGC**GGCAGCGTGCAGCTCGCC<br>**GCGGCGGCGCCACCACCA**GTCCTCGATGTTGTGGCGGATCTTGAAGTT |
| EGFP$^{R3(4)W3}$ | **CGCCGCCGCTGGTGGTGG**GGCAGCGTGCAGCTCGCC<br>**CCACCACCAGCGGCGGCG**GATGTTGTGGCGGATCTTGAAGTT |
| PTP1B$^{1W}$ | **TGGTGGTGGCGTCGACGTCGC**AATGACTATATCAACGCTAGTTTGATAAAAATGGAAGAAGC<br>**GCGACGTCGACGCCACCACCA**TTGATGTAGTTTAATCCGACTATGGTCAAAGGGAC |
| PTP1B$^{1R}$ | **CGTCGACGCCGTTGGTGGTGG**AATGACTATATCAACGCTAGTTTGATAAAAATGGAAGAAGC<br>**CCACCACCAACGGCGTCGACG**TTGATGTAGTTTAATCCGACTATGGTCAAAGGGACTG |
| PTP1B$^{2W}$ | **TGGTGGTGGCGTCGACGTCGC**AAAGAGATGATCTTTGAAGACACAAATTTGAAATTAAC<br>**GCGACGTCGACGCCACCACCA**TTTTTGTGGCCAGTATTGTGCGCA |
| PTP1B$^{2R}$ | **CGTCGACGCCGTTGGTGGTGG**AAAGAGATGATCTTTGAAGACACAAATTTGAAATTAACATTG<br>**CCACCACCAACGGCGTCGACG**TTTTTGTGGCCAGTATTGTGCGCATTTTAACG |
| PTP1B$^{3W}$ | **GGTTGGTGGTGGCGTCGACGCCGTGGC**ACCCAAGAAACTCGAGAGATCTTACATTTCCAC<br>**GCCACGGCGTCGACGCCACCACCAACC**TGTAAGGTTTTCCAATTCTAGCTGTCGCACTG |
| PTP1B$^{3R}$ | **GGTCGTCGACGCCGTTGGTGGTGGGGC**ACCCAAGAAACTCGAGAGATCTTACATTTCCAC<br>**GCCCCACCACCAACGGCGTCGACGACC**TGTAAGGTTTTCCAATTCTAGCTGTCGCACTG |
| PTP1B$^{4W}$ | **TGGTGGTGGCGTCGACGTCGC**CACGGGCCCGTTGTGGTGCAC<br>**GCGACGTCGACGCCACCACCA**CGGGCTGAGTGACCCTGACTCTC |
| PTP1B$^{4R}$ | CGTCGACGCCGTTGGTGGTGGCACGGGCCCGTTGTGGTGCA<br>**CCACCACCAACGGCGTCGACG**CGGGCTGAGTGACCCTGACTCTCGGACTTTGAAAAGAAAG |
| PTP1B$^{5W}$ | **GGTTGGTGGTGGCGTCGACGTCGC**GCCCAAAGGAGTTACATTCTTACCCAGG<br>**GCGACGTCGACGCCACCACCAACC**CATTTTTATCAAACTAGCGTTGATATAGTCATTATCTTCTTG |
| PTP1B$^{5R}$ | **GGTCGTCGACGCCGTTGGTGGTGG**GCCCAAAGGAGTTACATTCTTACCCAGG<br>**CCACCACCAACGGCGTCGACGACC**CATTTTTATCAAACTAGCGTTGATATAGTCATTATCTTCTTG |
| PNP$^{1R}$ | **CGTCGCCGACGTTGGTGGTGG**CACCGACCTCAAGTTGCAATAATC<br>**CCACCACCAACGTCGGCGACG**CTTAGTATGAGACAGAAGCCATTCTGCAG |
| PNP$^{2R}$ | **CGTCGCCGACGTTGGTGGTGG**GGCAGGGCCTGTGTGATGATGCA<br>**CCACCACCAACGTCGGCGACG**ATTCAGGAACCCAAACACCAGT |
| PNP$^{3R}$ | **CGTCGCCGACGTTGGTGGTGG**CAACGTGAGCTACAGGAAGGCA<br>**CCACCACCAACGTCGGCGACG**CCCCATTTGTTTCCAGGTACTGAGAG |

*Nucleotide sequences encoding inserted CPPs are shown in boldfaced letters.

**Table S2.** Amino Acid Sequence of Selected Proteins

| **Protein** | **Amino Acid Sequence** |
|---|---|
| **EGFP**$^{WT}$ | MDSLEFIASKLVSKGEELFTGVVPILVELDGDVNGHKFSVSGEGEGDATYGKLTLKFIC<br>TTGKLPVPWPTLVTTLTYGVQCFSRYPDHMKQHDFFKSAMPEGYVQERTIFFKDDG<br>NYKTRAEVKFEGDTLVNRIELKGIDFKEDGNILGHKLEYNYNSHNVYIMADKQKNGI<br>KVNFKIRHNIEDGSVQLADHYQQNTPIGDGPVLLPDNHYLSTQSALSKDPNEKRDH<br>MVLLEFVTAAGITLGMDELYKLEHHHHHH |
| **EGFP**$^{W3R3}$ | MDSLEFIASKLVSKGEELFTGVVPILVELDGDVNGHKFSVSGEGEGDATYGKLTLKFIC<br>TTGKLPVPWPTLVTTLTYGVQCFSRYPDHMKQHDFFKSAMPEGYVQERTIFFKDDG<br>NYKTRAEVKFEGDTLVNRIELKGIDFKEDGNILGHKLEYNYNSHNVYIMADKQKNGI<br>KVNFKIRHNIED**WWWRRR**GSVQLADHYQQNTPIGDGPVLLPDNHYLSTQSALSK<br>DPNEKRDHMVLLEFVTAAGITLGMDELYKLEHHHHHH |
| **EGFP**$^{R3W3}$ | MDSLEFIASKLVSKGEELFTGVVPILVELDGDVNGHKFSVSGEGEGDATYGKLTLKFIC<br>TTGKLPVPWPTLVTTLTYGVQCFSRYPDHMKQHDFFKSAMPEGYVQERTIFFKDDG<br>NYKTRAEVKFEGDTLVNRIELKGIDFKEDGNILGHKLEYNYNSHNVYIMADKQKNGI<br>KVNFKIRHNI**RRRWWW**GSVQLADHYQQNTPIGDGPVLLPDNHYLSTQSALSKDP<br>NEKRDHMVLLEFVTAAGITLGMDELYKLEHHHHHH |
| **EGFP**$^{R4W3}$ | MDSLEFIASKLVSKGEELFTGVVPILVELDGDVNGHKFSVSGEGEGDATYGKLTLKFIC<br>TTGKLPVPWPTLVTTLTYGVQCFSRYPDHMKQHDFFKSAMPEGYVQERTIFFKDDG<br>NYKTRAEVKFEGDTLVNRIELKGIDFKEDGNILGHKLEYNYNSHNVYIMADKQKNGI<br>KVNFKIRHNI**RRRRWWW**GSVQLADHYQQNTPIGDGPVLLPDNHYLSTQSALSKD<br>PNEKRDHMVLLEFVTAAGITLGMDELYKLEHHHHHH |
| **PTP1B**$^{WT}$ | MEMEKEFEQIDKSGSWAAIYQDIRHEASDFPCRVAKLPKNKNRNRYRDVSPFDHSRI<br>KLHQEDNDYINASLIKMEEAQRSYILTQGPLPNTCGHFWEMVWEQKSRGVVMLNR<br>VMEKGSLKCAQYWPQKEEKEMIFEDTNLKLTLISEDIKSYYTVRQLELENLTTQETRE<br>ILHFHYTTWPDFGVPESPASFLNFLFKVRESGSLSPEHGPVVVHCSAGIGRSGTFCLA<br>DTCLLLMDKRKDPSSVDIKKVLLEMRKFRMGLIQTADQLRFSYLAVIEGAKFIMGDS<br>SVQDQWKELSHEDLEPPPEHIPPPPRPPKRILEPHNVDSLEFIASKLAAALEHHHHH<br>H |
| **PTP1B**$^{1W}$ | MEMEKEFEQIDKSGSWAAIYQDIRHEASDFPCRVAKLPKNKNRNRYRDVSPFDHSRI<br>KLHQ**WWWRRR**RNDYINASLIKMEEAQRSYILTQGPLPNTCGHFWEMVWEQKSR<br>GVVMLNRVMEKGSLKCAQYWPQKEEKEMIFEDTNLKLTLISEDIKSYYTVRQLELEN<br>LTTQETREILHFHYTTWPDFGVPESPASFLNFLFKVRESGSLSPEHGPVVVHCSAGIG<br>RSGTFCLADTCLLLMDKRKDPSSVDIKKVLLEMRKFRMGLIQTADQLRFSYLAVIEG<br>AKFIMGDSSVQDQWKELSHEDLEPPPEHIPPPPRPPKRILEPHNVDSLEFIASKLAAA<br>LEHHHHHH |
| **PTP1B**$^{1R}$ | MEMEKEFEQIDKSGSWAAIYQDIRHEASDFPCRVAKLPKNKNRNRYRDVSPFDHSRI<br>KLHQ**RRRRWWW**NDYINASLIKMEEAQRSYILTQGPLPNTCGHFWEMVWEQKSR<br>GVVMLNRVMEKGSLKCAQYWPQKEEKEMIFEDTNLKLTLISEDIKSYYTVRQLELEN<br>LTTQETREILHFHYTTWPDFGVPESPASFLNFLFKVRESGSLSPEHGPVVVHCSAGIG<br>RSGTFCLADTCLLLMDKRKDPSSVDIKKVLLEMRKFRMGLIQTADQLRFSYLAVIEG<br>AKFIMGDSSVQDQWKELSHEDLEPPPEHIPPPPRPPKRILEPHNVDSLEFIASKLAAA<br>LEHHHHHH |

| | |
|---|---|
| **PTP1B$^{2R}$** | MEMEKEFEQIDKSGSWAAIYQDIRHEASDFPCRVAKLPKNKNRNRYRDVSPFDHSRI<br>KLHQEDNDYINASLIKMEEAQRSYILTQGPLPNTCGHFWEMVWEQKSRGVVMLNR<br>VMEKGSLKCAQYWPQK**RRRRWWW**KEMIFEDTNLKLTLISEDIKSYYTVRQLELEN<br>LTTQETREILHFHYTTWPDFGVPESPASFLNFLFKVRESGSLSPEHGPVVVHCSAGIG<br>RSGTFCLADTCLLLMDKRKDPSSVDIKKVLLEMRKFRMGLIQTADQLRFSYLAVIEG<br>AKFIMGDSSVQDQWKELSHEDLEPPPEHIPPPPRPPKRILEPHNVDSLEFIASKLAAA<br>LEHHHHHH |
| **PTP1B$^{4R}$** | MEMEKEFEQIDKSGSWAAIYQDIRHEASDFPCRVAKLPKNKNRNRYRDVSPFDHSRI<br>KLHQEDNDYINASLIKMEEAQRSYILTQGPLPNTCGHFWEMVWEQKSRGVVMLNR<br>VMEKGSLKCAQYWPQKEEKEMIFEDTNLKLTLISEDIKSYYTVRQLELENLTTQETRE<br>ILHFHYTTWPDFGVPESPASFLNFLFKVRESGSLSP**RRRRWWW**HGPVVVHCSAGIG<br>RSGTFCLADTCLLLMDKRKDPSSVDIKKVLLEMRKFRMGLIQTADQLRFSYLAVIEG<br>AKFIMGDSSVQDQWKELSHEDLEPPPEHIPPPPRPPKRILEPHNVDSLEFIASKLAAA<br>LEHHHHHH |
| **PNP$^{WT}$** | MRGSHHHHHHGMASMTGGQQMGRDLYDDDDKDPTLMENGYTYEDYKNTAEW<br>LLSHTKHRPQVAIICGSGLGGLTDKLTQAQIFDYSEIPNFPRSTVPGHAGRLVFGFLNG<br>RACVMMQGRFHMYEGYPLWKVTFPVRVFHLLGVDTLVVTNAAGGLNPKFEVGDI<br>MLIRDHINLPGFSGQNPLRGPNDERFGDRFPAMSDAYDRTMRQRALSTWKQMGE<br>QRELQEGTYVMVAGPSFETVAECRVLQKLGADAVGMSTVPEVIVARHCGLRVFGFSL<br>ITNKVIMDYESLEKANHEEVLAAGKQAAQKLEQFVSILMASIPLPDKAS |
| **PNP$^{3R}$** | MRGSHHHHHHGMASMTGGQQMGRDLYDDDDKDPTLMENGYTYEDYKNTAEW<br>LLSHTKHRPQVAIICGSGLGGLTDKLTQAQIFDYSEIPNFPRSTVPGHAGRLVFGFLNG<br>RACVMMQGRFHMYEGYPLWKVTFPVRVFHLLGVDTLVVTNAAGGLNPKFEVGDI<br>MLIRDHINLPGFSGQNPLRGPNDERFGDRFPAMSDAYDRTMRQRALSTWKQMG**R**<br>**RRRWWW**QRELQEGTYVMVAGPSFETVAECRVLQKLGADAVGMSTVPEVIVARHC<br>GLRVFGFSLITNKVIMDYESLEKANHEEVLAAGKQAAQKLEQFVSILMASIPLPDKAS |

*Inserted CPP sequences are shown in boldfaced letters.

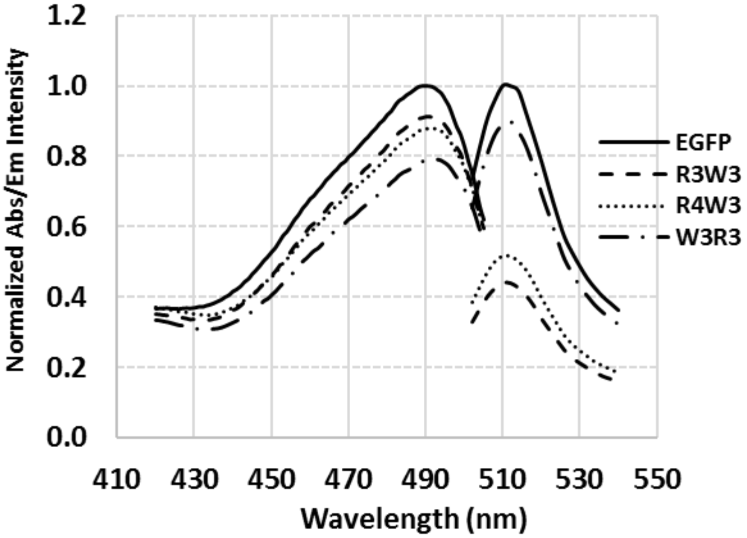


**Figure S1.** Comparison of the absorption (420-505 nm) and emission spectra (500-540 nm) of wild-type and mutant EGFP.

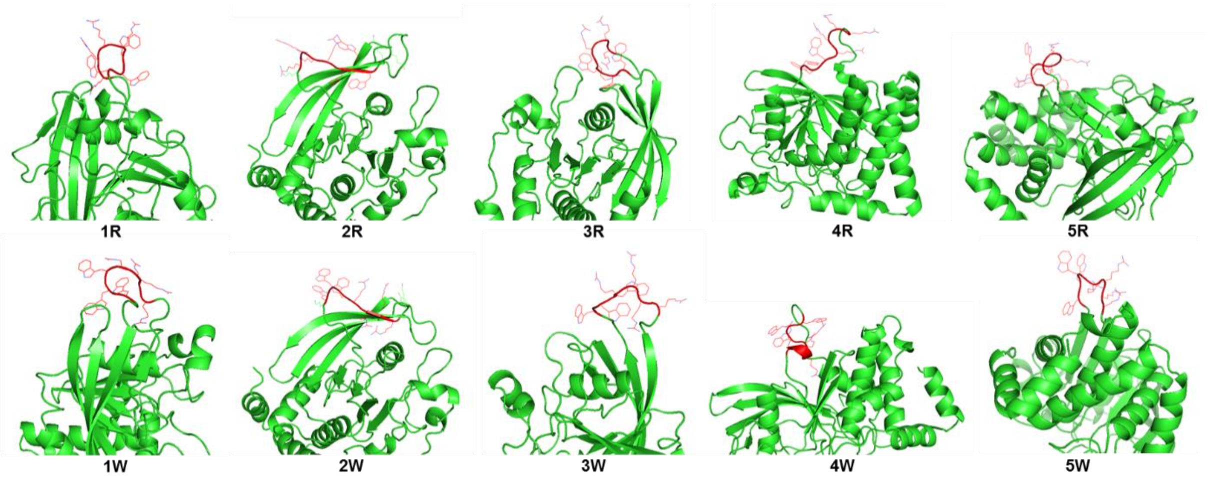


**Figure S2.** Predicted protein folds of PTP1B loop insertion mutants. CPP sequences are highlighted in red with side chain depicted. Structures were analyzed by PyMOL.

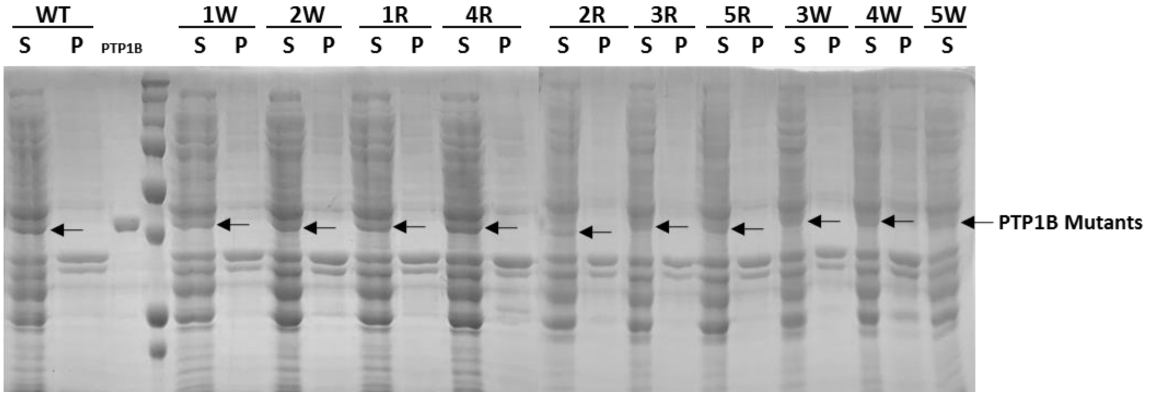


**Figure S3.** SDS-PAGE showing the pilot-scale expression of 10 PTP1B mutants (5 mL of culture). S, soluble fraction of the cell lysate; P, insoluble fraction of the cell lysate.

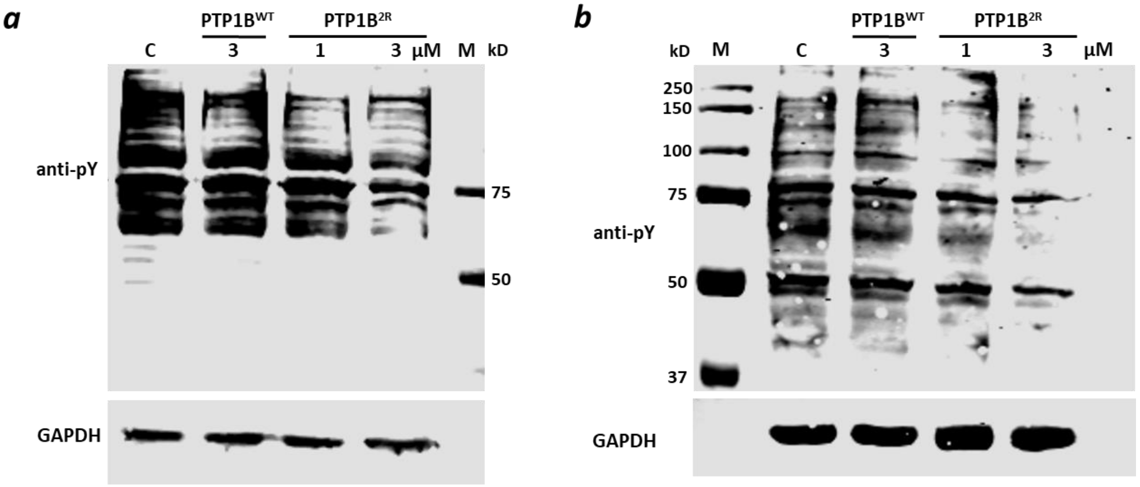


**Figure S4.** Effect of PTP1B$^{WT}$ and PTP1B$^{2R}$ (1 or 3 μM) on the global pY levels in HeLa (*a*) and A549 cells (*b*). C, control cells without PTP1B treatment.

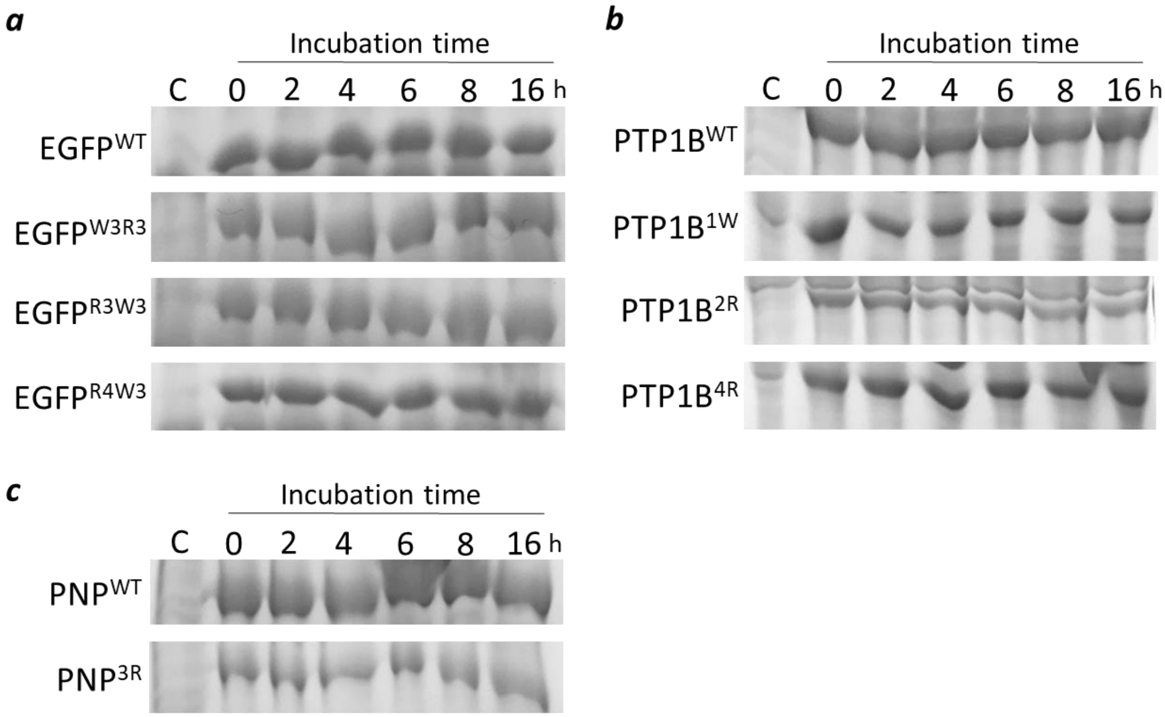


**Figure S5.** Coomassie blue stained SDS-PAGE gel (10%) showing the degradation of wild-type and mutant EGFP (*a*), PTP1B (*b*), and PNP proteins (*c*) by serum proteases.

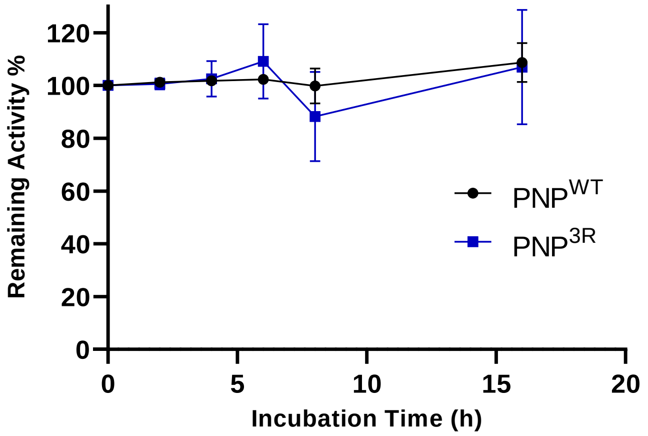


**Figure S6.** Serum stability of wild-type and mutant PNP as monitored by quantitating the remaining enzymatic activities after varying periods of incubation.

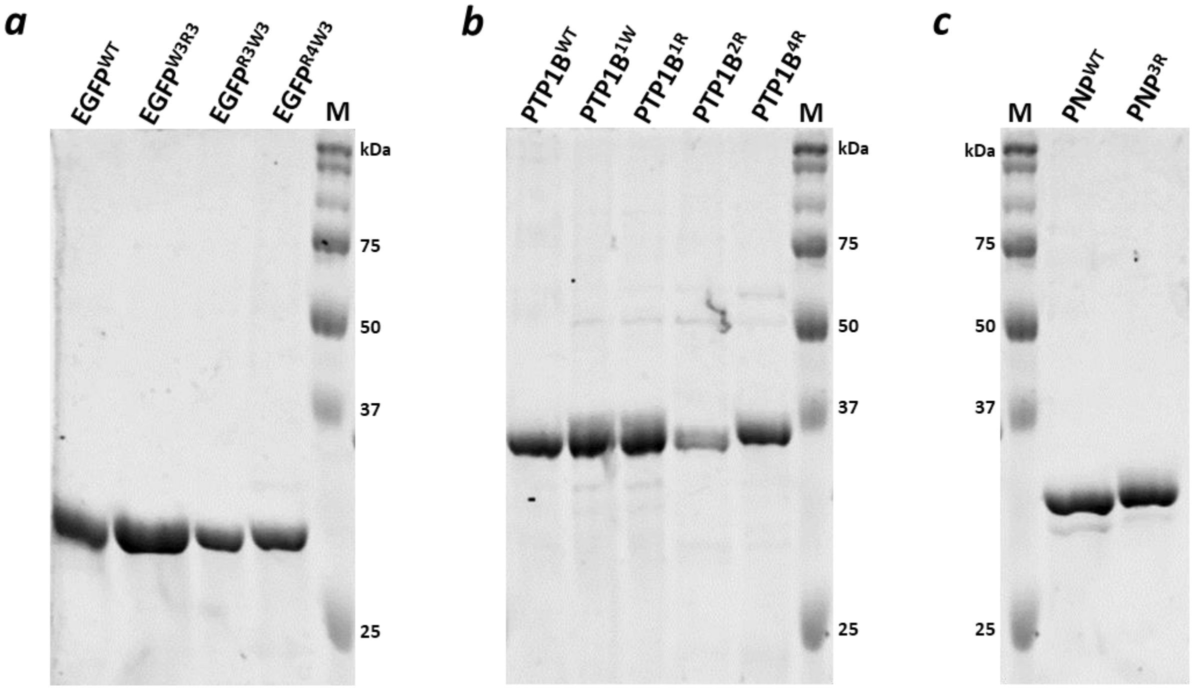


**Figure S7.** Coomassie blue stained SDS-PAGE showing the purities of wild-type and mutant EGFP (*a*), PTP1B (*b*), and PNP protein samples (*c*) used in this work. M, molecular weight markers.
